# Modelling the brain response to arbitrary visual stimulation patterns for a flexible high-speed BCI

**DOI:** 10.1101/358036

**Authors:** Sebastian Nagel, Martin Spüler

**Author notes:** Corresponding author *Email address* (Sebastian Nagel).

## Abstract

Visual evoked potentials (VEPs) can be measured in the EEG as response to a visual stimulus. Commonly, VEPs are displayed by averaging multiple responses to a certain stimulus or a classifier is trained to identify the response to a certain stimulus. While the traditional approach is limited to a set of predefined stimulation patterns, we present a method that models the general process of VEP generation and thereby can be used to predict arbitrary visual stimulation patterns from EEG and predict how the brain responds to arbitrary stimulation patterns. We demonstrate how this method can be used to model single-flash VEPs, steady state VEPs (SSVEPs) or VEPs to complex stimulation patterns. It is further shown that this method can also be used in a BCI to allow information transfer rates of more than 470 bit/min and lead to more flexible BCIs with a virtually unlimited amount of targets and any desired trial duration.

## 1. Introduction

Visual evoked potentials (VEPs) are electrical potentials that can be measured by electroencephalography (EEG) as the brain’s responses to a visual stimulus. VEPs are used in a variety of fields ranging from clinical diagnostics (Chirapapaisan et al., 2015) over basic research (Blake & Logothetis, 2002) to their application in a Brain-Computer Interface (BCI) (Wolpaw et al., 2002).

The idea of a VEP-based BCI was originally developed by Sutter (1984), who proposed ”the visual evoked response as a communication channel” and envisioned the use of VEPs for a BCI-controlled keyboard. Sutter implemented such a VEP-based BCI in 1992 where he used 64 visual stimuli that were modulated by a complex stimulation pattern.

Today, VEP-based BCIs are either based on the original idea of Sutter and use complex stimulation patterns (also called codes) to elicit a code-modulated VEP (cVEP) (Spüler et al., 2012; Bin et al., 2011), or they use visual stimulation with a specific frequency which evokes steady-state VEPs (SSVEPs) (Chen et al., 2015). Although both methods differ in how the stimulation pattern is constructed, both methods depend on the construction of a stimulus-specific template, restricting the number of possible targets.

One notable exception is the work by Thielen et al. (2015), who investigated a more general approach to use visual stimulation patterns that are not restricted to a predefined pattern. Their model is based on the assumption that the response to a complex stimulation pattern is a linear superposition of individual single-flash VEPs (Capilla et al., 2011). Thielen et al. therefore developed a convolution model that breaks down a complex stimulation patterns in smaller subcomponents and models the response to a complex stimulation pattern by a superposition of the responses to the subcomponents. While this approach allows for a more flexible prediction model, the stimulation patterns are not fully arbitrary, as they can only be composed of short and long pulses.

It should be noted, that the usability of stimulation patterns is restricted by the brain. Herrmann (2001) has shown that brain responses can only be found by using stimulation frequencies of up to 90 Hz. Furthermore, the stimulation rate is limited by the used hardware for stimulus presentation, like a computer monitor. Therefore, ”arbitrary stimulation patterns” should be interpreted as a huge set of possible stimulation patterns relative to the used stimulation rate.

In this paper, we present a model which is able to predict arbitrary stimulation patterns and its backward model which is able to predict the brain response to arbitrary stimulation patterns. We show that both models can be used for high-speed BCI control using any desired trial duration with a virtually unlimited amount of targets.

## 2. Methods

### 2.1. EEG2Code model

#### 2.1.1. Training

The model is based on a ridge regression model, which is able to interpret the EEG signal and predict the stimulation pattern during an arbitrary stimulation. For training, the stimulation pattern is always a fully random binary sequence presented with a rate of 60 bit/s. Since the most prominent parts (N1, P1 and N2) of a VEP response to a single stimulus lasts for around 250 ms, we use a 250 ms window of the spatially filtered EEG data as predictor and the corresponding bit of the stimulation pattern as response to train the ridge regression model. The window is shifted sample-wise over the data during a trial, meaning that it is required to use 250 ms of EEG data after trial end, otherwise the last 250 ms of a stimulation pattern can not be predicted. Fig. 1(A) depicts this procedure with a bit-wise window shifting for simplicity. The ridge regression model 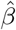 and its bias term *β*_0_ can be calculated by

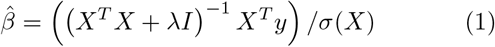

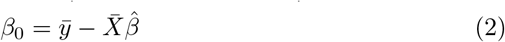

where *X* (the predictor) is a *n × k*-matrix with *n* the number of windows and *k* the window length (number of samples). *y* (the responses) is a *n ×* 1-vector containing the corresponding bit of the modulation sequence for each window. *I* is the identity matrix and *λ* the ridge regression parameter, which was not optimized but set to 0.001. Since a window has a length of *k* =150 samples, at the used sampling rate *s* =600*Hz*, the output 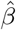 is a coefficient vector of length 150, one for each input sample and the constant bias term *β*_0_. The number of windows *n* depends on the number of trials *N*, the average trial duration *d*, the window length *k* and the sampling rate *s*:

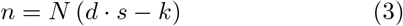

As described in section 2.5.4, we used *N* =96 and *d* =4*s* resulting in *n* =216, 000 windows.

**Figure 1:**
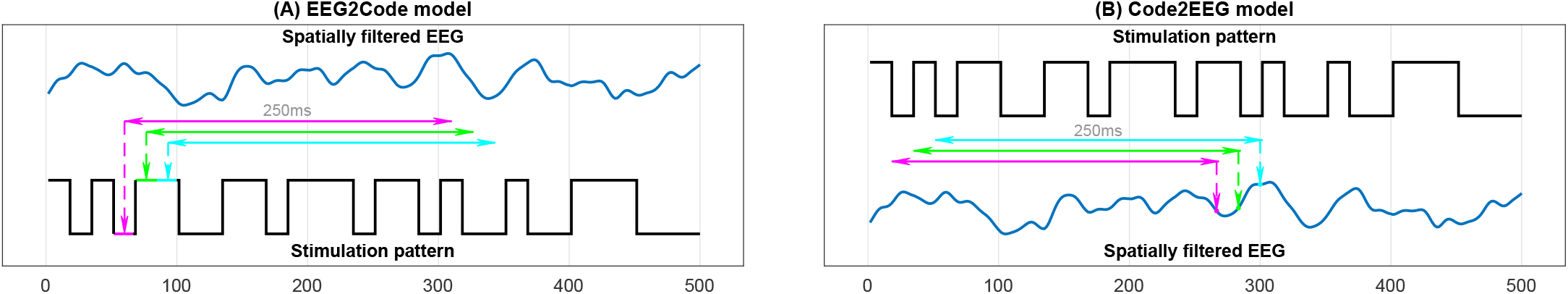
**(A)** Training of the EEG2Code model. Each 250ms window of the spatially filtered EEG data will be projected to its corresponding bit (1 or 0) of the corresponding stimulation pattern. **(B)** Training of the Code2EEG model. Each 250ms window of the modulation pattern will be projected to the corresponding value of the spatially filtered EEG data.

#### 2.1.2. Prediction

After training, the model is able to predict the bits of the stimulation pattern. Fig. 2(B) depicts the procedure, the measured EEG data is spatially filtered and the trained regression model is used to predict each 250 ms window (sample-wise shifted). The regression model predicts a real number *yi* for each window *i*

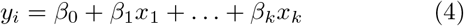

where *β*_0_ is the constant term and *β*_1_*…k* the coefficients for each window sample *x_k_*. The predicted real values *y_i_* in turn can be interpreted as a binary values by a simple threshold method, each value above or equal 0.5 is set to be binary 1 and 0 otherwise. Afterwards, the predicted binary sequence is compared to the stimulation pattern using the hamming distance. The binary transformation is only done for identifying the bit prediction accuracy, not for BCI control (see section 2.3).

**Figure 2:**
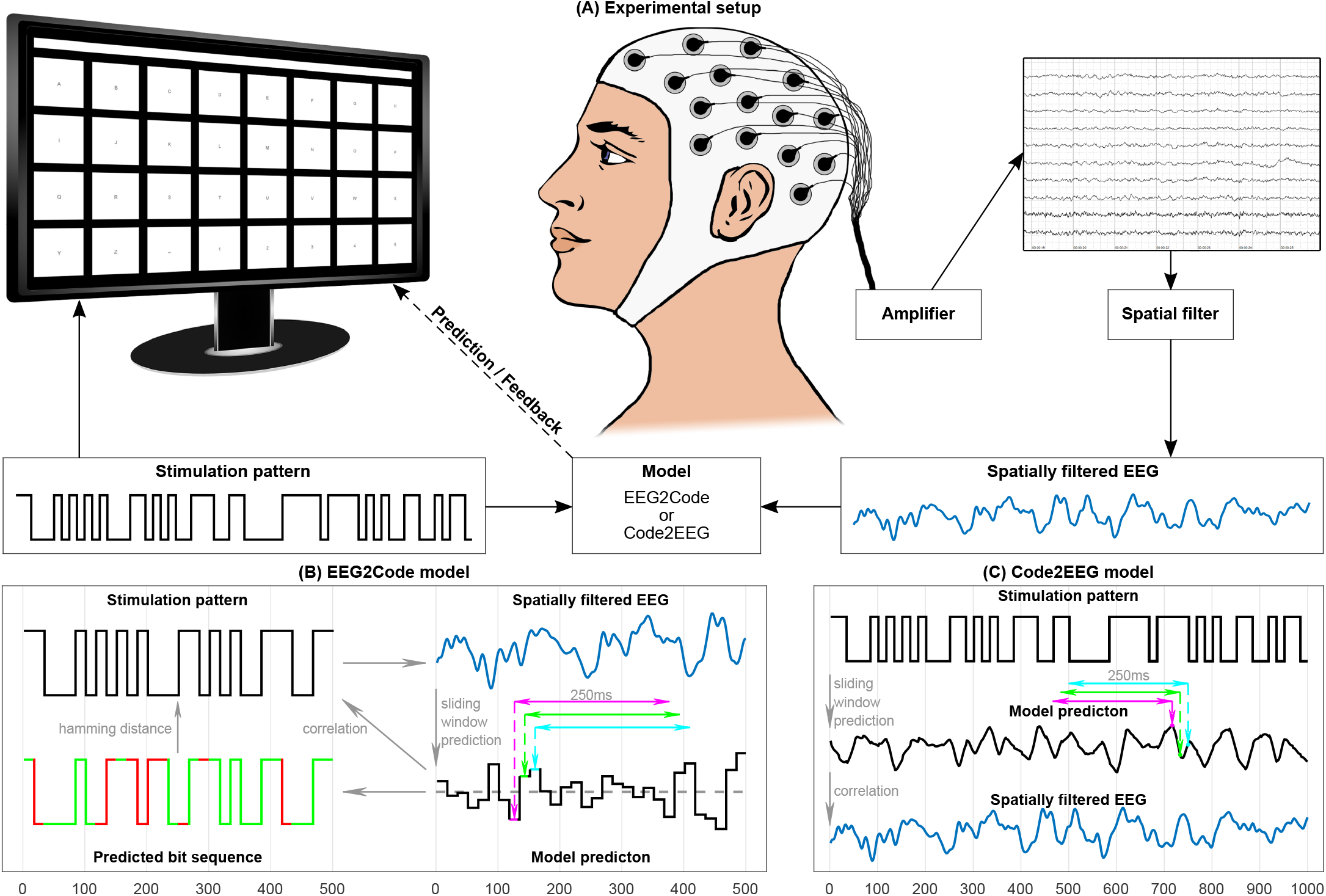
**(A)** Setup of the BCI experiment. The presentation layout is as shown on the monitor, it has 32 targets labeled alphabetically from A to Z followed by ‘-’ and numbers 1 to 5. The targets are separated by a blank black space and above targets is the text field showing the written text during the experiment. Each target is modulated with its own random stimulation pattern. During a trial, the participant has to focus a target. A spatial filter is applied to the measured EEG. The model predicts the target which is indicated to the participant by highlighting the target in yellow and the letter is appended to the text field above the keyboard. **(B)** The EEG2Code model predicts an arbitrary stimulation pattern. A 250ms window will be slided sample-wise over the spatially filtered EEG signal. For simplicity, it is shown bit-wise using 3 exemplary windows. The trained model predicts a real number for each window. Each value above 0.5 (gray dashed line) is interpreted as boolean 1 or 0, otherwise. The resulting bit sequence can be compared (hamming distance) to the stimulation pattern (match =green, mismatch =red). For target selection we used the correlation coefficient between model prediction and the stimulation patterns of all targets, the one which correlates most will be chosen as the predicted target. **(C)** The Code2EEG model predicts the brains response to an arbitrary stimulation pattern by sliding the 250 ms window over the stimulation pattern. For BCI control, the model predicts the brain responses to the stimulation patterns of all targets, which in turn are compared to the measured EEG using the correlation coefficient. The one which correlates most will be chosen as the predicted target.

### 2.2. Code2EEG model

#### 2.2.1. Training

Like the EEG2Code model, the Code2EEG model is also based on a ridge regression model and trained on fully random stimulation patterns to predict the EEG response based on a stimulation pattern. We also use 250 ms windows (150 samples) and the equation is the same as Eqn. 1 and 2, but the predictors *X* are the windows of the stimulation pattern and the response *y* is the spatially filtered EEG data. Fig. 1(B) depicts this procedure for three exemplary windows.

Again, the output 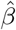 of the ridge regression is a coefficient vector of length 150. And we set *λ* to 0.001.

#### 2.2.2. Prediction

After training, the model is able to predict the brain response to an arbitrary stimulation pattern. The equation is the same as Eqn. 4, but *x*_1_ *… x_k_* are the *k* samples of the modulation pattern window. The model prediction *y* can now be compared to the measured (and spatial filtered) EEG. We used the correlation coefficient to compare them. Fig. 2(C) depicts the procedure, in this case, the real stimulation pattern is used to predict the brain response.

#### 2.2.3. Modelling brain response

As described, the model can predict the brain response to an arbitrary stimulation pattern, therefore, we also analyzed the prediction of a single flash stimulus, a 30 Hz SSVEP stimulus, and the m-sequence used for spatial filter generation (see section 2.6). For this, an additional participant (not included in the BCI experiment) had to perform the spatial filter and training session followed by 120 single flash trials and 120 SSVEP trials using 30 Hz stimulation frequency, whereas each trial lasts for 1 s. We predicted the brain response to those stimuli and compared them to the spatially filtered EEG data.

### 2.3. BCI control

For BCI control, each target is modulated with its own (random) stimulation sequence and we need a method to choose the correct target out of others. For the EEG2Code model we used the correlation coefficient between the model prediction and the modulation patterns of all targets. The corresponding target which correlates most is selected. We could also use the pattern prediction accuracy, but as shown in our previous study (Nagel et al., 2017), this is leading to a loss of additional information which in turn leads to a reduced performance.

The Code2EEG model predicts the EEG data of the modulation patterns of all targets and compares them to the measured EEG data, again, the one which correlates most is chosen as the correct target.

### 2.4. Modulation patterns

#### 2.4.1. Random modulation patterns

During the experiment the MT19937 (Matsumoto & Nishimura, 1998) random generator was used for generating random modulation patterns. At each monitor refresh a random integer (0 or 1) is generated for each target, therefore, the binary sequence of a target is always random without conscious repetitions and generated with a rate of 60 bit/s, continuously. The pattern generation can be repeated or varied by using the same or a different random seed, respectively.

#### 2.4.2. Optimized modulation patterns

In our previous study (Nagel et al., 2018b) we found that the number of bit changes is a crucial property of stimulation patterns that leads to different performances. We analyzed the pattern prediction accuracy of all 250 ms (15 bit) sub-sequences and found a maximum accuracy for sub-sequences with 7 bit changes. We repeated the analysis using the data of the current study and confirmed the findings. The pattern prediction accuracy of 15 bit sub sequences is best for sequences with 6 to 8 bit changes (Fig. 3), meaning a bit change probability of approximately 50% at a stimulation rate of 60 Hz.

**Figure 3:**
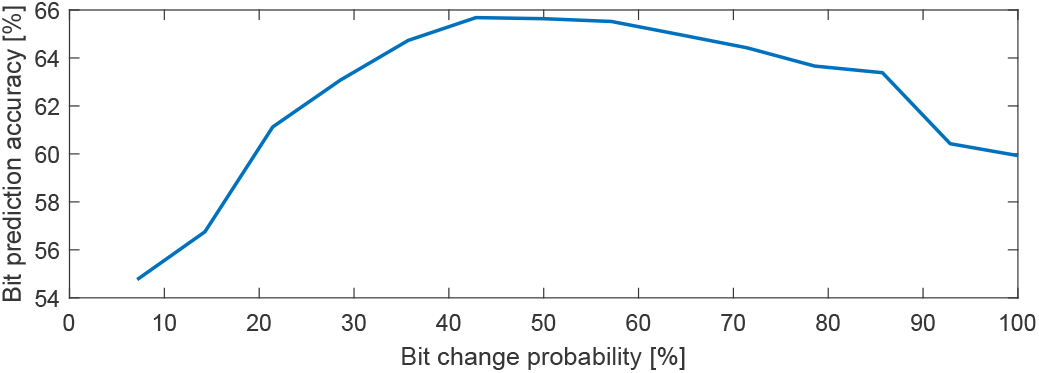
Average bit prediction accuracy of the EEG2Code model of all 250 ms (15 bit) sub-sequences of random modulation trials grouped by the bit change probability. For example, a bit change probability of 100% means that each successive bit changes from 0 to 1 or 1 to 0, respectively. The maximum bit prediction accuracy is reached between approximately 40% to 60%, meaning an average of 6 to 8 bit changes. Therefore, we used only 15 bit subsequences with 7 bit changes for the optimized modulation patterns.

Therefore, we generated a set of 15 bit long sequences with 7 bit changes, this results in a total number of 6,864 bit sequences. As we use a correlation measure to determine the correct target, we filtered those sequences. For this, we generated 100,000 subsets of 150 randomly chosen sequences out of the 6,864 bit sequences and took the subset with lowest average correlation between the sequences in the subset. The resultant subset has an average correlation of −0.004 (SD =0.276) between any sub-sequence to all others. The subset allows to modulate 150^*T*/250ms^ different targets, where *T* is the trial duration in milliseconds.

### 2.5. Experimental setup

#### 2.5.1. Hardware & Software

The BCI consists of a g.USBamp (g.tec, Austria) EEG amplifier, two personal computers (PCs), Brainproducts Acticap system with 32 channels and a LCD monitor (BenQ XL2430-B) for stimuli presentation. Participants are seated approximately 80 cm in front of the monitor.

PC1 is used for the presentation on the LCD monitor, which is set to refresh rate of 60 Hz and its native resolution of 1920 × 1080 pixels. A stimulus can either be black or white, which can be represented by 0 or 1 in a binary sequence and is synchronized with the refresh rate of the LCD monitor. The timings of the monitor refresh cycles are synchronized with the EEG amplifier by using the parallel port.

PC2 is used for data acquisition and analysis. As a general framework for recording the data of the EEG amplifier we used BCI2000 (Schalk et al., 2004) and the data processing is done with MATLAB (2017). The amplifier sampling rate was set to 600 Hz, resulting in 10 samples per frame/stimulus. Additionally, a TCP network connection was established to PC1 in order to send instructions to the presentation layer and to get the modulation patterns of the presented stimuli.

We used a 32 electrodes layout, 30 were located at Fz, T7, C3, Cz, C4, T8, CP3, CPz, CP4, P5, P3, P1, Pz, P2, P4, P6, PO9, PO7, PO3, POz, PO4, PO8, PO10, O1, POO1, POO2, O2, OI1h, OI2h, and Iz. The remaining two electrodes were used for electrooculography (EOG), one between the eyes and one left of the left eye. The ground electrode (GND) was positioned at FCz and reference electrode (REF) at OZ.

#### 2.5.2. Presentation layout

We used MATLAB (2017) and the Psychtoolbox (Brainard & Vision, 1997) for the presentation layer. Our layout is a 4 × 8 matrix keyboard layout (32 targets in total) as shown in Fig. 2(A), whereas the targets are labeled alphabetically from A to Z followed by ‘_’ and numbers 1 to 5. The targets are separated by a blank black space and above targets is a text field showing the written text.

#### 2.5.3. Participants

The study was approved by the local ethics committee of the Medical Faculty at the University of Tübingen and conformed to the guidelines of the Declaration of Helsinki. A written informed consent was obtained from all participants. To test the system, 9 healthy subjects (5 female) were recruited. All subjects had normal or correctedto-normal vision. The age ranged from 18 to 23 years. Each subject participated in one session and completed the whole experiment. None of the subjects participated in other VEP EEG studies before.

#### 2.5.4. Data acquisition

Initially, the participants had to perform a run to generate a spatial filter (see section 2.6.3). The training phase was split into 3 runs, but with varying trial duration, 5 s, 4 s, and 3 s, respectively. The testing phase was split into 14 runs with a trial duration of 2 s. The runs were alternated using random stimulation patterns and optimized stimulation patterns.

During all runs the inter-trial time was set to 0.75 s and the participants had to perform 32 trials in lexicographic order (see section 2.5.2).

### 2.6. Preprocessing

#### 2.6.1. Frequency filter

The recorded EEG data is bandpass filtered by the amplifier between 0.1 Hz and 60 Hz using a Chebyshev filter of order 8 and an additional 50 Hz notch filter was applied.

#### 2.6.2. Correcting raster latencies

Standard computer monitors (CRT, LCD) cause raster latencies because of the line by line image build-up dependent on the refresh rate. As VEPs are effected by these latencies resulting in a decreased BCI performance, we corrected the raster latencies as described in our previous work (Nagel et al., 2018a).

#### 2.6.3. Spatial filter

Recent studies (Bin et al., 2011; Spüler et al., 2012) have shown increased classification accuracy by using spatial filters to improve the signal-to-noise ratio of the brain signals. As random stimulation is not suitable for spatial filter training, a m-sequence with low auto-correlation is used for target modulation. The spatial filter training is done using a canonical correlation analysis (CCA) as described in a previous work (Spüler et al., 2014), except that the presentation layout and stimulation duration differ. The presentation layout is as described above and the participants had to perform 32 trials (one per target) whereas one trial lasts for 3.15 seconds followed by 1.05 for gaze shifting. As the used modulation pattern has a length of 63 bits (1.05 seconds), we got 96 sequence cycles per participant, which in turn are used for training the spatial filter. The spatial filter is then used for the following experiment.

### 2.7. Performance evaluation

For the pattern prediction of the EEG2Code model, the binary values of the predicted patterns were compared to the stimulation patterns by using the hamming distance. Because the distances are 1’s and 0’s, the averaged hamming distance of all samples corresponds to the accuracy of how much of the stimulation pattern can be predicted correctly.

The prediction of the brain responses is evaluated by using the correlation coefficient between the predicted EEG and the spatially filtered EEG.

The BCI control performance of both models is evaluated using the accuracy of correctly predicted targets.

Additional to the accuracies, we calculated the corresponding information transfer rates (ITRs) (Wolpaw et al., 1998). The ITR can be computed with the following equation:

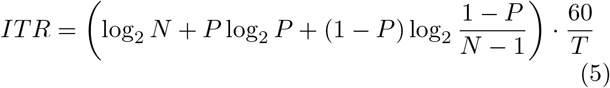

with *N* the number of classes, *P* the accuracy, and *T* the time in seconds required for one prediction. The ITR is given in bits per minute (bpm).

For the pattern prediction of the EEG2Code model the values are: *N* =2 and *T* = ^1^*/*60 s. For BCI control *N* equals the number of targets and *T* the trial duration including the inter-trial time.

## 3. Results

To summarize, the method presented in this paper allows to create prediction models in two directions: on the one hand the EEG2Code model can be used to predict the visual stimulation pattern based on the EEG, on the other hand the Code2EEG model can be used to predict the brain response (the VEP measured by EEG) for a given visual stimulation pattern. Both models can be used for BCI control, for which multiple stimuli (i.e. targets) are modulated with random patterns, as depicted in Figure 2. For data acquisition we performed an online BCI control experiment, additionally we did an offline analysis to get the performance of the stimulation pattern prediction and the brain response prediction. Furthermore, we analyzed the BCI performance by varying the trial duration and the number of targets. The results are shown in the following subsections.

### 3.1. Stimulation pattern prediction

When using random visual stimulation, the EEG2Code model is able to predict the stimulation pattern with an average accuracy of 64.6% (the percentage of correctly predicted bits in the bit-vector), which corresponds to an ITR of 232 bit per minute (bpm). For the best subject, the model is able to predict 69.1% of the stimulation pattern correctly, which corresponds to an ITR of 389.9 bpm.

### 3.2. Brain response prediction

Using the Code2EEG model, we can predict the EEG response to a visual stimulation pattern. Exemplary, the model trained on random visual stimulation was used to predict the response to a single-light flash of 16.6 ms, to a 30 Hz SSVEP pattern, and to a cVEP m-sequence pattern (Fig. 4). When predicting the EEG response to random visual stimulation patterns, the average correlation between the prediction and measured EEG is r=0.346, with a maximum correlation of r=0.465 for subject S5 (see Table 1 for details).

**Figure 4:**
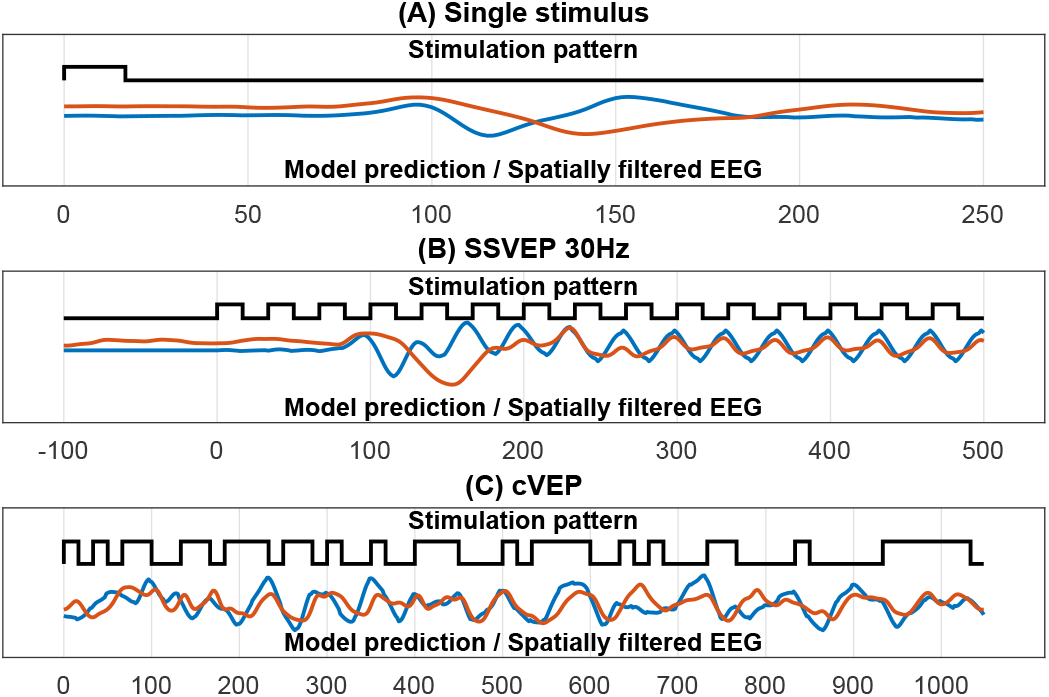
Predicted brain responses of the Code2EEG model compared to the measured EEG (120 trials averaged). The black line represents the stimulation pattern, the blue line the predicted brain response and the red line the spatially filtered EEG. **(A)** Single stimulus pattern lasting for 1/60 s. **(B)** 30 Hz SSVEP pattern. **(C)** cVEP pattern.

**Table 1:** Results of the model prediction. Accuracy (ACC) and information transfer rates (ITR) of the EEG2Code model, and correlation (r) of the Code2EEG model prediction to the measured EEG data. Shown are the average results of all subjects. Best results are in bold font.

| Subject | EEG2Code |  | Code2EEG<br>r |
| --- | --- | --- | --- |
|  | ACC [%] | ITR [bpm] |  |
| S1 | **69.1** | **389.9** | 0.425 |
| S2 | 64.5 | 222.4 | 0.240 |
| S3 | 63.7 | 196.5 | 0.351 |
| S4 | 65.6 | 257.8 | 0.394 |
| S5 | 66.3 | 282.7 | **0.465** |
| S6 | 67.1 | 308.2 | 0.403 |
| S7 | 63.7 | 196.8 | 0.226 |
| S8 | 60.9 | 124.5 | 0.324 |
| S9 | 60.2 | 109.4 | 0.287 |
| mean | 64.6 | 232.0 | 0.346 |

### 3.3. Brain-Computer Interface control

As mentioned, both models can be used for BCI control. In the following we show the online performance followed by the offline analysis by varying the trial duration and the number of targets.

#### 3.3.1. Online results

The EEG2Code model was tested in an online BCI with a trial length of 2 s. An inter-trial time of 0.75 s was chosen, because previous experiments have shown that 0.5 s is too short for untrained users. The participants, who never used a BCI before, had to perform 7 runs with fully random stimulation patterns and 7 runs with optimized stimulation patterns. During each run the participants had to spell each letter in lexicographic order, meaning 32 trials per run for a total of 224 trials. Table 2 shows the target prediction accuracies and the corresponding ITRs. When using completely random stimulation patterns the average accuracy of target selection is 97.8%, which corresponds to 103.9 bpm. As we found that modulation patterns with a specific number of bit changes lead to better results (Nagel et al., 2017), optimized modulation sequences (details in Material and Methods section) were also tested, leading to an accuracy of 98.0% (108.1 bpm). Due to a ceiling effect, the difference between random and optimized stimulation patterns is not significant.

**Table 2:** Results of the online BCI experiment. Accuracies (ACC) and information transfer rates (ITR) of the online BCI experiment (EEG2Code model). Shown are the average results of all subjects using random stimulation patterns and optimized stimulation patterns. ITRs are calculated including the inter-trial time of 0.75s.

| Subject | optimized |  | random |  |
| --- | --- | --- | --- | --- |
|  | ACC [%] | ITR [bpm] | ACC [%] | ITR [bpm] |
| S1 | 100.0 | 109.1 | 100.0 | 109.1 |
| S2 | 99.6 | 107.7 | 99.1 | 106.5 |
| S3 | 99.1 | 106.5 | 96.4 | 100.4 |
| S4 | 100.0 | 109.1 | 100.0 | 109.1 |
| S5 | 100.0 | 109.1 | 99.1 | 106.5 |
| S6 | 100.0 | 109.1 | 100.0 | 109.1 |
| S7 | 99.6 | 107.7 | 98.2 | 104.3 |
| S8 | 99.1 | 106.5 | 96.9 | 101.3 |
| S9 | 84.8 | 79.3 | 90.6 | 89.2 |
| mean | 98.0 | 108.1 | 97.8 | 103.9 |

#### 3.3.2. Varying trial duration and number of targets

Both models are based on a sliding window approach and are trained on fully random stimulation patterns, therefore, the trial duration can be varied and it is not required to use the same stimulation patterns for training and testing, but to use arbitrary stimulation patterns which in turn allows to vary the number of targets. The following results are based on the runs with optimized stimulation patterns.

Using the EEG2Code model with 32 targets, varying the trials length from 0.5 s to 2 s shows that average accuracy increases with longer trials, but the ITR reaches its optimum of 154.3 bpm with a trial duration of 1 s. It is worth mentioning that S1 achieved an ITR of 231.1 bpm using 600 ms trial duration which corresponds to an accuracy of 92.41%. The Code2EEG model is generally less accurate, reaching only 94.4 % with 2 s trials, but still has an optimum average ITR of 146.6 bpm with a trial length of 0.75 s. More detailed results for the EEG2Code model are shown in Fig. 5(A) and (B) and for the Code2EEG model in Fig. A1(A) and (B).

**Figure 5:**
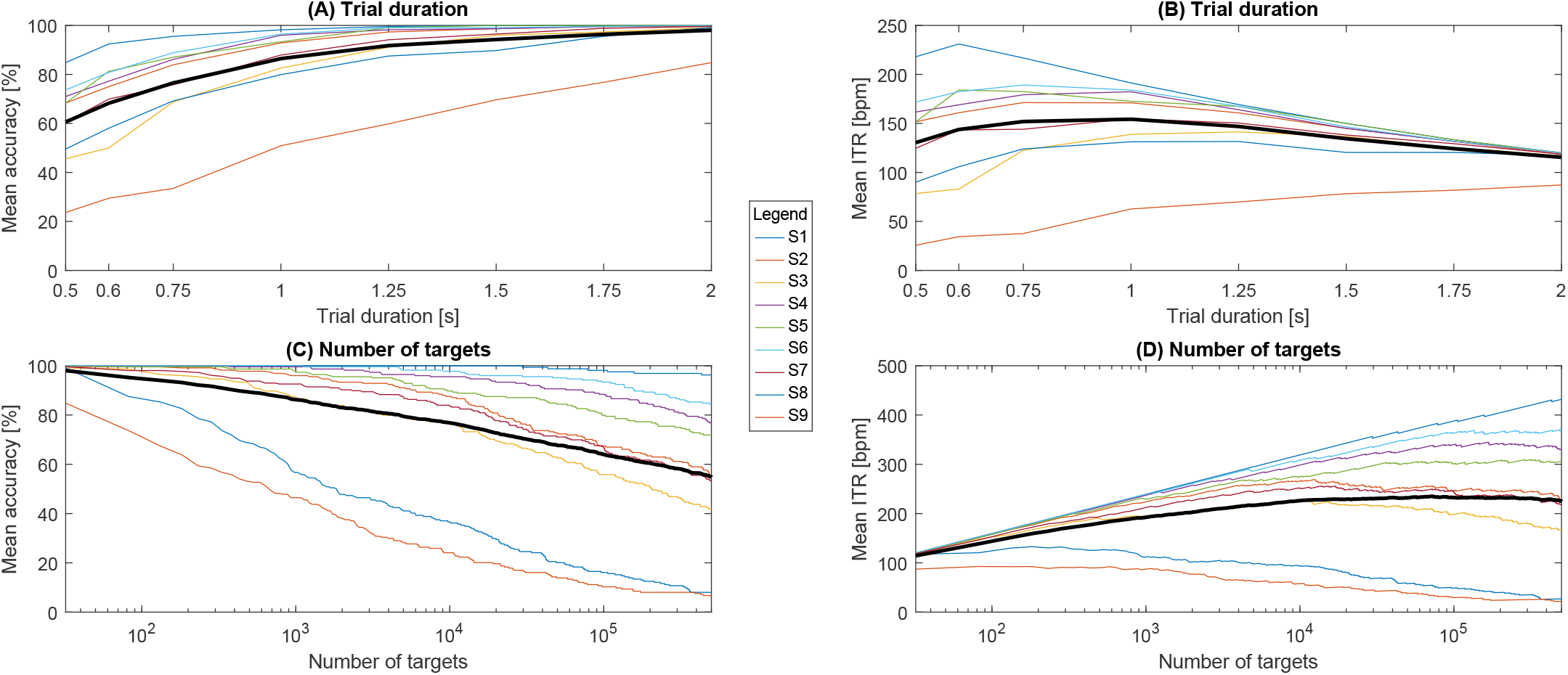
Results using the EEG2Code model. Shown are the accuracies and ITRs (including 0.5 s inter-trial time). (A) and (B) using varying trial durations and (C) and (D) using varying number of targets (logarithmic scale). Each colored line is one subject and the thick black line represents the mean of all subjects.

As both models are not limited to fixed stimulation patterns, but work with arbitrary patterns, the number of targets can be increased. With a trial length of 2 s and 60 Hz refresh rate, there are 2^120^ = 1.33 *·* 10^36^ different stimulation patterns, which therefore is the upper bound for the number of targets.

For this paper, we simulated a BCI with up to 500,000 targets in steps of 50 targets. Fig. 5(C) and (D) shows the accuracies and ITRs of the EEG2Code model relative to the number of targets. The averaged optimum ITR was at 235.3 bpm using 71,930 targets, although the optimum varied largely between subjects. For subject S1 the ITR was still increasing at half a million targets with an an accuracy of 96.3% and an ITR of 432 bpm. Furthermore, S1 achieved an accuracy of 100% for up to 29,500 targets. In Fig. A2 are detailed results for S1 showing the target variation for all trial durations, revealing a maximum ITR of 474.5 bpm using 472,700 targets and a trial duration of 1.5 s, which is known to us, the highest reported ITR of an offline BCI.

Using the Code2EEG model, the maximum average ITR of 183.8 bpm is reached with 9,600 targets with an average accuracy of 64.93%. Adding more targets decreases the ITR for most of the participants. Detail results for the Code2EEG model can be found Fig. A1(C) and (D).

## 4. Discussion

In this paper, we presented a method that models the process of VEP generation and can be used in two directions to either predict the EEG response to a visual stimulation pattern (Code2EEG) or predict the visual stimulation pattern from the EEG (EEG2Code). Contrary to previous methods (Thielen et al., 2015; Cardona et al., 2016), the presented method works with arbitrary stimulation patterns, while it is only trained on a limited set of stimulation patterns. We used random stimulation patterns because we assume to cover most of the possible VEP responses.

Using the EEG2Code model, we have demonstrated that a stimulation pattern presented at 60 Hz can be predicted with an average accuracy of 64.6 %, which corresponds to an ITR of 232 bpm. It should be noted, that a binary classification method, like a support vector machine, could lead to better stimulation pattern prediction, but for comparison we used the continuous regression method (with threshold) as we also used the regression model for brain response prediction, which has to be continuous.

When using the Code2EEG model to predict the EEG response to a certain stimulation pattern, the correlation between the recorded EEG and the predicted EEG response is *r* =0.346. It should be noted that the Code2EEG model only predicts the evoked response in the EEG, and not the noise present in the EEG, so that the correlation between the prediction and the evoked response is higher. On the other side, a correlation of *r* =0.346 means that *r*^2^=11.97% of the variance in the recorded EEG can be explained by the Code2EEG model, thereby giving a lower bound for the signal-to-noise-ratio (SNR) of the visual evoked response in the EEG. This is confirmed by comparing the model prediction of the cVEP pattern to the averaged recorded EEG (Fig.4(C)), this resulted in a correlation of *r* =0.551.

As the presented approach is based on the assumption of linearity in the VEP generation process (Capilla et al., 2011; Lalor et al., 2006), we have shown that most of the VEP response to arbitrary stimulation patterns can be explained by a superposition of single VEP responses. But interestingly, the duration of the predicted single-flash VEP response and the onset of the predicted 30 Hz SSVEP response is shortened in time compared to the recorded VEP responses, whereas the sinusoidal part of the 30 Hz SSVEP response matches roughly perfect. It seems there is an initially slowed response after a longer stimulation pause which the model can not reflect because it is trained using random stimulation patterns where longer stimulation pauses are unlikely. By using a model trained on sequences with *<* 3 bitchanges the onset matches better for all stimulation patterns, whereas the further prediction matches worse (Fig. A3). This could be an evidence, that the VEP generation is a non-linear process. Furthermore, this could be the reason for the worse stimulation pattern prediction of sequences with less bit changes (Fig. 3).

We have also shown that both models can be used for controlling a BCI. When using the EEG2Code model for an online BCI, we achieved an average accuracy of 108 bpm and thereby, with an ITR >100 bpm, fall in the category of high-speed BCIs. But the performance of our online EEG2Code BCI is below the best results reported for a cVEP BCI (144 bpm, Spüler et al., 2012) or an SSVEP BCI (267 bpm, Chen et al., 2015). However, the BCI used in the online setup was not optimized to achieve a high ITR, but to be comfortably usable by a BCI-naive person and to get the required data for the offline analysis. In the offline analysis we have shown that the parameters can be optimized, for example by reducing the trial duration, to achieve an average ITR of 154.3 bpm and up to 231.1 bpm. For the best subject, we found a theoretical maximum ITR of 474.5 bpm, although we only simulated results for up to 500,000 targets and the ITR was still rising at that point, so that it is likely that the maximum is even higher.

Compared to other types of BCIs, the high number of possible targets is a unique feature of the EEG2Code BCI. With 1000 targets, the average accuracy is still around 90 % and goes down to around 55 % for half a million targets, with the best subject still achieving >95 % accuracy. As the *Dictionary of Chinese Variant Form* compiled by the Taiwan (ROC) Ministry of Education in 2004 contains 106,230 individual characters, this BCI approach would theoretically allow to select each character of that alphabet individually. Although this thought is purely theoretical as there are practical limitations, like displaying such a high amount of targets.

The Code2EEG model resulted in a lower BCI performance compared to the EEG2Code model, but with a trial length of 0.75 s it achieved an average ITR of 146.5 bpm, which is three times the ITR as the comparable re-convolution BBVEP of Thielen et al. (2015), which achieved an average ITR of 48.4 bpm.

Since our methods are based on a sliding window approach allowing to vary the trial duration, they can easily be used for an asynchronous BCI, for example by only classifying if a specific correlation is reached.

Due to the discussed assumption of non-linearity, we expect a better pattern prediction of the EEG2Code model using a non-linear method. Achieving accuracies of always above 75% would allow to use error-correction codes, as know from coding theory. This in turn would allow to encode arbitrary information as a binary sequence, directly transfer it through visual stimuli, decode it and correct possible errors. This would allow to build more robust and fully flexible BCI applications by using the brain as data transfer channel.

Finally, the approach to create a general model for the generation of VEPs could also be applied to other stimulation paradigms like sensory or auditory stimulation. Also, such an approach could be utilized for electrical stimulation for therapeutic means, where optimal stimulation parameters need to be found to evoke a certain response (Walter et al., 2014).

## Acknowledgements

This work was supported by the *Deutsche Forschungsgemeinschaft* (DFG; grant SP-1533*\* 2-1).

## Author contributions statement

S.N. and M.S. conceived and designed the study and wrote the paper; S.N. performed the experiment and analyzed the data.

## Additional information

The authors declare no conflict of interest.

